# Crosstalk between Proline Hydroxylation and Histidine Methylation Regulates Actin-Dependent Cell Mobility Control

**DOI:** 10.1101/2021.08.26.456853

**Authors:** Zhengshuai Liu, Zhenzhen Zi, Chiho Kim, Xu-dong Wang, Yonghao Yu

## Abstract

Actin filaments generate intrinsic forces that are essential for cell motility. This process is tightly regulated by various posttranslational modifications (PTMs), including acetylation, arginylation, and oxidation. However, the role of actin hydroxylation in regulating its dynamics remains poorly understood. Here, we demonstrate that the inhibition of Prolyl Hydroxylase Domain-containing proteins (PHDs) activity significantly promotes actin polymerization and enhances cell motility. Using hydroxylation proteomics, we identified actin as a substrate for PHDs, with hydroxylation occurring at multiple proline residues, especially at proline 70 (Pro70). This modification recruits the von Hippel-Lindau (VHL) tumor suppressor protein, which then leads to the disruption of the interaction between actin and the histidine methyltransferase SET Domain Containing 3 (SETD3). Consequently, SETD3-mediated methylation of histidine 73 (His73) on actin is suppressed, impairing actin polymerization and compromising cell motility. Notably, genetic loss of VHL or pharmacological inhibition of PHDs restores His73 methylation, enhances actin filament formation, and promotes cell motility. Together, our findings uncover a novel regulatory crosstalk between hydroxylation and methylation on actin, establishing a critical mechanism by which PTMs fine-tune actin dynamics to govern cell motility.

**Highlights:**

- MK-8617 augments cell motility and actin polymerization in VHL dependent manner;
- Hydroxylation proteomics reveals that actin is hydroxylated by PHDs;
- Hydroxylation at Pro70 recruits VHL, disrupting actin’s interaction with SETD3 and reducing His73 methylation;
- Genetic loss of VHL in clear cell renal cell carcinoma (ccRCC) potentially promotes cell motility by increasing His73 methylation on actin.

## Introduction

Actin is a highly conserved cytoskeletal protein that plays a fundamental role in cell motility, shape maintenance, and structural integrity^1^. Within cells, actin exists in two distinct forms: monomeric (G-actin) and filamentous (F-actin). The dynamic polymerization of G-actin into F-actin forms filaments that provide essential intracellular mechanical support, central to its cytoskeletal function^1, 2^. Actin dynamics are tightly regulated by post-translational modifications (PTMs) that fine-tune actin’s polymerization and interactions with other binding partners^3, 4^. Notably, certain PTMs, such as phosphorylation at Tyr53^5^, ADP-ribosylation at Arg177^6^, and S-glutathionylation at Cys374^7^ are known to inhibit actin polymerization. In contrast, other PTMs like Nt-arginylation^8^ and methylation at His73^9^ enhance actin polymerization. Specially, the His73 methylation of actin has been reported to be catalyzed by the methyltransferase, SET domain protein 3 (SETD3), a PTM event that plays a critical role in regulating smooth muscle contractility^9^. Emerging research suggests that novel types of PTMs may occur on actin, yet the full spectrum of these modifications and their potential roles in regulating actin dynamics remain largely uncharted. Besides, understanding whether distinct PTMs on actin interact synergistically or antagonistically could reveal key mechanisms by which actin is precisely regulated in response to cellular signals and environmental changes.

Hydroxylation is a PTM that influences protein stability, interactions, and functional activity. This modification occurs on various amino acids, including proline, lysine, asparagine, aspartate and histidine^10^. The proline hydroxylation reaction is catalyzed by the 2-oxoglutarate- and oxygen-dependent dioxygenases Prolyl Hydroxylase Domain-containing enzymes PHD1, PHD2, and PHD3^11^. Hypoxia-inducible factor alpha (HIFα) is a well-established substrate hydroxylated by PHDs. Prolyl-hydroxylated HIFα is recognized by the von Hippel–Lindau protein (pVHL)–elongin C (ELC)–elongin B–cullin 2 (CUL2)–RBX1 (VCB–CR) E3 ubiquitin ligase complex and is targeted for ubiquitylation and proteasomal degradation^12^. Notably, a large proportion of renal cancers, especially clear cell renal cell carcinoma (ccRCC), are marked by the loss of VHL tumor suppressor function, resulting in the accumulation of hypoxia-inducible factors, particularly HIF-2α, a key driver of ccRCC^13^. Beyond VHL recognition and proteasomal degradation, the potential for non-ubiquitin-dependent pathways regulating protein function following proline hydroxylation remains largely unexplored. Uncovering these novel mechanisms will pave the way for the development of innovative and potent therapies for kidney cancer.

Our study demonstrated that MK-8617, a pan-inhibitor of PHD enzymes, significantly enhanced actin polymerization and cell motility in a VHL-dependent manner. Subsequent hydroxylation proteomic analyses further identified proline hydroxylation on actin, illuminating the molecular mechanism by which hydroxylation affects actin dynamics. Specifically, we found that (1) Pro70 on actin can be hydroxylated by PHD enzymes; (2) Pro70 hydroxylation recruits VHL, which then inhibits the interaction between actin and SETD3; and (3) this disruption reduces His73 methylation on actin, which in turn inhibits actin polymerization and impairs cell motility. In conclusion, our findings reveal that actin hydroxylation modulates actin dynamics through methylation, offering new insights for the development of therapies targeting actin-related diseases.

## Materials and Methods

### Reagents and antibodies

MK-8617 was purchased from Selleckchem (S8443), and was used at 5 μM final concentration; TRITC– conjugated phalloidin was purchased from (P1951), and was used at 1 µg/ml. Isopropyl-β-Dthiogalactopyranoside (IPTG) (Dioxane-free) was obtained from Fisher Scientific (BP1755). β-Actin (13E5) Rabbit mAb was obtained from Cell Signaling (4970s). Anti-H73Me antibody was kindly provided by Dr. Or Gozani (Stanford University School of Medicine); Additional reagents were purchases from commercial vendors, including Anti-GAPDH (Thermo Fisher Scientific, AM43000), HIF-1α Antibody (Cell Signaling, 3716s), VHL antibody (Cell Signaling, 68547s), SETD3 Antibody (Bethyl laboratories, A304-072A), PHD-2/Egln1 (D31E11) Rabbit mAb (Cell Signaling, 4835s), EGLN3/PHD3 Antibody (Novus Biologicals, NB100-139), Anti-CUL-2 Antibody (C-4) (Santa Cruz, sc-166506), Anti-Flag (Sigma, F7425), Anti-Flag (Sigma, F1804), Anti-HA (Cell Signaling Technology, 3724s), Donkey anti Rabbit IgG (GE healthcare, NA9340), Goat anti Mouse IgG (Millipore, AP181P), Monoclonal Anti-HA−Agarose (Sigma Aldrich, A-2095), ANTI-FLAG M2 Affinity Gel (Sigma Aldrich, A2220), GST (26H1) Mouse mAb (Cell Signaling, 2624s) and HRP-Conjugated Streptavidin (Thermo Fisher, 21130).

### Cell Culture

Hela (ATCC), and HEK293T cells were cultured in Dulbecco’s modified Eagle’s medium (DMEM, sigma, D5796) supplemented with 10% fetal bovine serum (FBS, sigma, 12306C) and antibiotics (Gibco, 15240112) at 37 °C in a 5% CO2 incubator. 786-O, and RCCJF cells were cultured in RPMI-1640 medium (sigma, R8758).

### Lentiviral packaging and stable cell line construction

HEK293T cells were cultured to 50–70% confluency in 10 cm dishes, and were then transfected using Lipofectamin-2000 (Sigma). The Plenti or pLKO.1 plasmid, VSVG, and delta8.9 were cotransfected at 8 μg, 6 μg, and 4 μg, respectively. The medium was changed 6 hours after the transfection. Viruses were collected twice at 24 h and 48 h after the transfection. Subsequently, 1 mL of virus was added to each well of Hela or 786-O cells in 6-well plates with Polybrene at a final concentration of 8 μg / ml. After splitting the cells once, cells were infected with virus again using the same procedure. The medium was replaced after 48 h with a fresh growth medium containing 2 μg/ml blasticidin or puromycin.

### Tandem Mass Tags (TMT) Mass Spectrometry

After the compound treatment, the cells were lysed using 1% SDS lysis buffer (1% SDS, 10 mM HEPES, pH 7.0, 2 mM MgCl_2_, 20 U/ml universal nuclease with added protease inhibitors and phosphatase inhibitors). Protein lysates were reduced by 2 mM dithiothreitol, and were alkylated by adding iodoacetamide to a final concentration of 50 mM, followed by incubation in the dark for 20 min. The lysates were extracted by methanol-chloroform-precipitation and the proteins were solubilized in 8 M urea, then digested with Lys-C. The samples were then diluted to a final concentration of 2 M urea by the addition of 100 mM ammonium bicarbonate (pH = 7.8). Proteins were digested overnight with sequencing-grade trypsin at a 1:100 (enzyme/substrate) ratio. Digestion was quenched by the addition of trifluoroacetic acid to a final concentration of 0.1%. For TMT MS samples, peptides were desalted using SepPak C18 columns (Waters) according to the manufacturer’s instructions. Peptides were fractionated by using an off-line two-dimensional SCX-RP-HPLC (strong-cation-exchange reversed-phase HPLC) protocol. The sample were desalted using stage tips and subjected to MS analysis.

Proteomic samples were analyzed using LC-MS/MS approach on an LTQ Velos Pro Orbitrap mass spectrometer (Thermo).Tandem mass spectra were searched against a composite database consisting of the human UniProt protein database and its reversed sequences using the Sequest algorithm embedded in an in-house-developed software suite^14^. Database search parameters included a static modification of carbamidomethylation on cysteine residues (+57.02146 Da), and dynamic modifications of methionine oxidation (+15.99491 Da) and proline hydroxylation (+15.99491 Da). Search results were filtered to achieve a peptide-level false discovery rate (FDR) of <1% by applying a linear discriminant analysis incorporating multiple scoring metrics including XCorr, deltaCN, number of missed cleavages, precursor charge state (excluding singly charged peptides), precursor mass accuracy, peptide length, and the fraction of matched fragment ions. Peptide quantification was performed using the CoreQuant algorithm implemented in an in-house-developed software suite^15^.

### Immunoblot Analysis

For immunoblot analysis, the cells were lysed using 1% SDS lysis buffer (1% SDS, 10 mM HEPES, pH = 7.0, 2 mM MgCl_2_, 20 U/ml universal nuclease with added protease inhibitors and phosphatase inhibitors). Protein concentrations were measured using the BCA assay (23227, Thermo Fisher). The lysates were mixed with the 4X reducing buffer (60 mM Tris-HCl, pH = 6.8, 25% glycerol, 2% SDS, 14.4 mM 2-mercaptoethanol, 0.1% bromophenol blue). Samples were boiled for 10 min and the same amounts of proteins were subjected to electrophoresis using the standard SDS-PAGE method. Proteins were then transferred to a 0.22 μm nitrocellulose membrane (GE, 10600001), and were blocked with a TBST buffer (25 mM Tris-HCl, pH = 7.5, 150 mM NaCl, 0.05% Tween-20) containing 3% non-fat dried milk and incubated overnight with primary antibodies (1:1,000 dilution) at 4 °C and for 1h at room temperature with peroxidase-conjugated secondary antibodies (1:10000 dilution). Blots were developed using enhanced chemiluminescence and were exposed on autoradiograph films and were developed using standard methods. The results were quantified using Image J and analyzed using Prism GraphPad Software.

### In Vitro Methylation Assay

In vitro methyltransferase reactions were performed with 5 μg SETD3, 10 μg substrate peptide (hydroxylated or non-hydroxylated peptide), with or without VHL protein and 80 μM SAM (Sadenosylmethionine) in a buffer that contained 50 mM Tris pH = 8.0, 20 mM KCl and 5 mM MgCl_2_ and were incubated at 30 °C overnight. Reactions were resolved by dot blot analysis.

### CO-Immunoprecipitation Assay

HEK293T cells were rinsed once with ice-cold PBS 24-36 hours after the transfection and were lysed in the ice-cold BLB lysis buffer (10 mM KPO_4_, pH = 7.6, 6 mM EDTA, pH = 8.0, 10 mM MgCl_2_, 0.5 % NP-40, 0.1 % BriJ-35, 0.1 % DOC, adjust pH to 7.4) or IP buffer (0.5% NP-40, 150 mM NaCl, 50 mM Tris-HCl, pH = 7.5) added with EDTA-free protease inhibitors (Roche). The soluble fractions from cell lysates were isolated by centrifugation at 13,000 rpm for 10 minutes. For immunoprecipitations, 20 μl of a 50% slurry of anti-FLAG (A2220, Sigma) or anti-HA beads (A2095, Sigma) were added to each lysate and incubated with rotation for overnight at 4°C. Immunoprecipitants were washed three times with BLB lysis buffer. Immunoprecipitated proteins were denatured by the addition of 100 μl of the sample buffer and boiling for 10 minutes and subjected to immunoblot analysis. For the peptide and VHL co-immunoprecipitation assay, peptides (the hydroxylated or nohydroxylated peptide) were incubated with streptavidin beads for 1 hour, and then were incubated with HEK293T cell lysates for 2 hours. After washing 3 times, the samples were subjected to western blot analysis.

### Immunostaining Assay

Cells were plated on micro cover glasses (Electron Microscopy Sciences 72226-01). After reaching confluence, cells were washed with PBS and were fixed with 4% paraformaldehyde (PFA) in PBS for 10 min. The cells were permeabilized (0.2% Triton X-100, 10 min), and were blocked in 3% BSA (Sigma, A9418). The cells were then incubated with Phalloidin– Tetramethylrhodamine B isothiocyanate (P1951) for one hour at room temperature. After the incubation with primary antibodies, slides were washed three times, and were then stained with DAPI (Sigma, D9542). The slides were sealed with the mounting medium (FluorSave reagent, Millipore, 345789). Photomicrographs were taken of representative 10×, 20× or 40× magnification fields using a confocal microscope (Confocal Zeiss LSM880 Airyscan).

### Cell Scratch Assay

Cells were seeded onto 6 well plates and were cultured overnight to reach 50% confluence. The cell layer was scratched with a sterile 1-ml pipette tip, labeled, and photographed at the indicated time points with a microscope.

### Statistical analysis

Data are expressed as mean±SEM. Statistical significance was evaluated using the unpaired two-tailed Student’s t-test for two groups. For more than two groups, data were analyzed by one-way ANOVA or two-way ANOVA. Differences were considered significant at a *P* value < 0.05.

## Results

### PHD inhibitors augment actin polymerization and cell motility in a VHL-dependent manner

Actin is a critical cytoskeletal protein that drives cellular motility, with its dynamics finely regulated by post-translational modifications (PTMs) such as phosphorylation, acetylation, and methylation. However, actin hydroxylation remains poorly characterized. To investigate this modification and its role in cell motility, we treated VHL-proficient HeLa and VHL-deficient 786-O and RCCJF cells with MK-8617, a pan-prolyl hydroxylase (PHD) inhibitor. Scratch assays revealed that MK-8617 significantly enhanced cell motility in Hela cells whereas this effect was markedly attenuated in 786-O and RCCJF cells (Fig. 1A and Fig. S1). Interestingly, consistent with the effects of MK-8617 treatment, VHL knockdown in HeLa cells similarly enhanced cell motility. In contrast, VHL reintroduction in 786-O cells significantly suppressed cell motility (Fig. 1B).

**Fig. 1.**
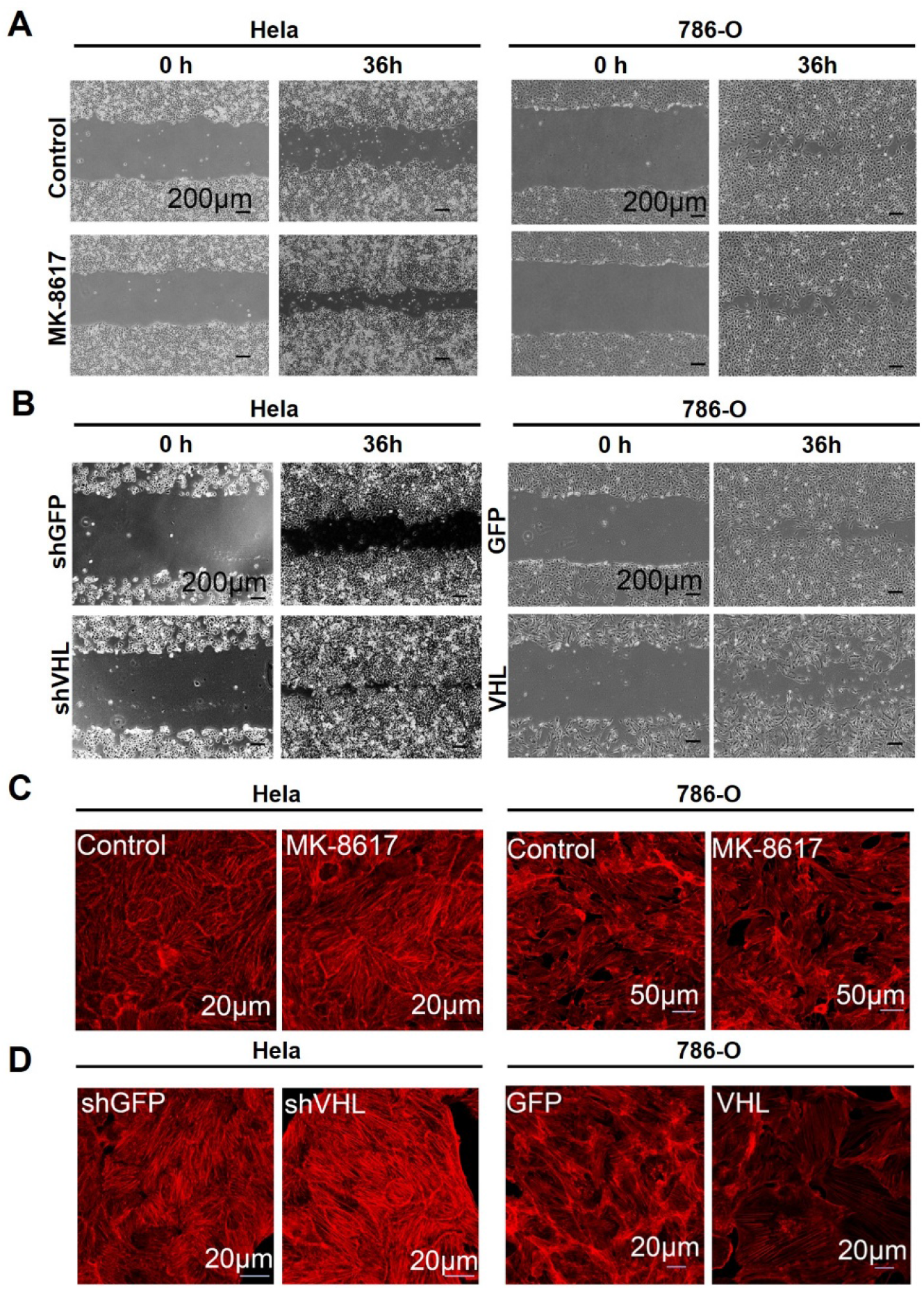
MK-8617 augments cell motility and actin polymerization in a VHL-dependent manner. A. MK-8617 promotes cell motility in Hela cells but not in 786-O cells. HeLa and 786-O cells were subjected to scratch assays with or without MK-8617 treatment (5 μM) for 36 hours. B. VHL knockdown increases cell motility in HeLa cells, whereas VHL overexpression decreases cell motility in 786-O cells. Scratch assays were performed on HeLa cells with or without VHL knockdown, and on 786-O cells with or without VHL overexpression. Representative images from two independent experiments were shown. Scale bar, 200 μm. C. MK-8617 increases β-actin polymerization in Hela cells but not in 786-O cells. HeLa and 786-O cells were treated with or without MK-8617 (5 μM) for 24 hours. D. VHL increases β-actin polymerization. VHL knockdown increases β-actin polymerization in Hela cells while VHL overexpress decrease β-actin polymerization in 786-O cells. Cells were fixed, permeabilized, stained with TRITC–conjugated phalloidin, and imaged by fluorescence microscopy. Scale bar, 20 μm or 50 μm.

To determine whether actin polymerization is influenced by MK-8617, we performed TRITC-conjugated phalloidin staining. The results showed that MK-8617 significantly increased actin polymerization in HeLa cells, whereas actin polymerization in 786-O cells exhibited insensitivity to MK-8617 treatment (Fig. 1C). Similarly, VHL knockdown in HeLa cells promoted actin polymerization, whereas VHL reintroduction in 786-O cells significantly suppressed actin polymerization (Fig. 1D). Collectively, these findings indicate that MK-8617 enhances actin polymerization and cell motility, likely via a mechanism that is dependent on the VHL protein.

### Hydroxy-Pro proteomic analyses identify actin as a Pro-hydroxylated protein

To determine whether PHDs mediate the hydroxylation of actin, we first assessed the interaction between PHD isoforms and actin using immunoprecipitation analysis. The results demonstrated that both PHD2 and PHD3 interact with actin (Fig. 2A). Next, we conducted TMT-labeled quantitative proteomic analysis to evaluate global hydroxylation changes in MCF10A cells following a time-course treatment with MK-8617 (Fig. 2B).

**Fig. 2.**
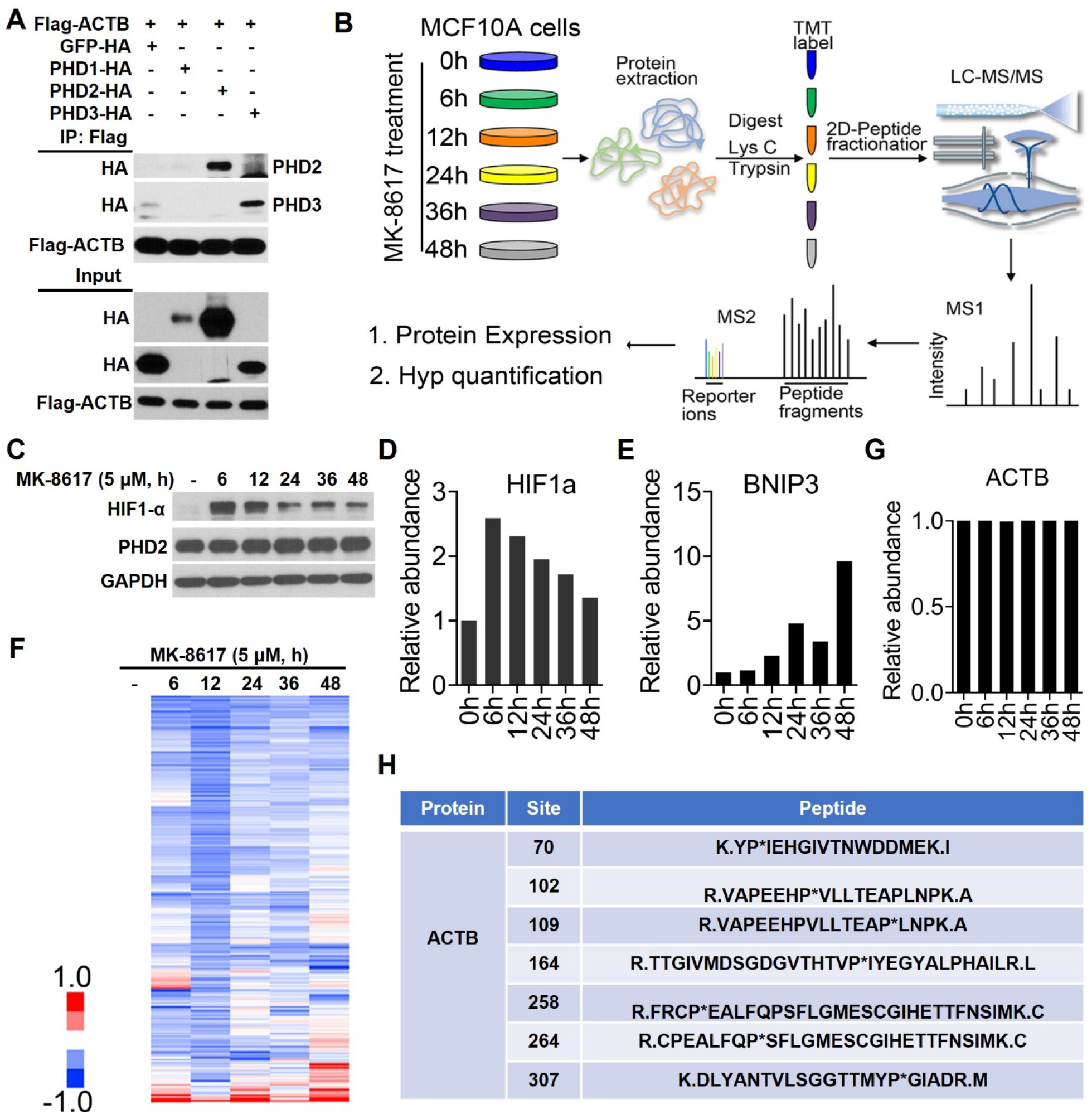
Hydroxy-proteomics identifies PHD-mediated hydroxylation of actin. A. Actin interacts with PHD2 and PHD3. Flag-tagged actin was co-transfected with HA tagged GFP, PHD1, PHD2 and PHD3 into HEK293T cells. The lysates were subjected to immunoprecipitation using the anti-Flag beads. PHD1-3 were probed using an antibody against the HA-tag. B. Overview of the workflow of the quantitative proteomic experiments. MCF10A cells were treated with MK-8617 (5 μM) upon indicated time points. Cell lysates were collected and subjected to TMT6-mass spectrometric analyses. C. HIF1α levels were measured using western blot upon MK-8617 (5 μM) treatment at the indicated time points. GAPDH was used as the internal control. (D-F). TMT-based quantitative proteomics results of HIF1a, BNIP3, and ACTB levels upon MK-8617 (5 μM) treatment. G. Global protein hydroxylation levels decreased following MK-8617 treatment. Hierarchical clustering of Pro-hydroxylated protein profiles in MCF10A cells treated with MK-8617 (5 μM) at the indicated time points. H. The hydroxy-Proline sites on actin and its corresponding peptide were identified through proteomics analyses.

Western blot analysis revealed that HIF-1α levels peaked at 6 hours post-MK-8617 treatment, followed by a gradual decline with extended exposure (Fig. 2C), a trend corroborated by the proteomic data (Fig. 2D). Similarly, BNIP3, a transcriptional target of HIF-1α, exhibited a time-dependent increase in protein expression in response to MK-8617 (Fig. 2E). In parallel, a significant downregulation of global protein hydroxylation was observed (Fig. 2F). These data validate the efficacy of MK-8617 treatment and confirm the reliability of the proteomic mass spectrometry data.

Subsequent analysis of actin expression and its hydroxylation status revealed that MK-8617 treatment did not affect total actin levels. Notably, multiple hydroxylated peptides were identified on actin, including proline residues at 70, 102, 109, 164, 258, 264, and 307 (Fig. 2G and Fig. 2H). These data demonstrate that actin could undergo hydroxylation mediated by PHDs.

### Hydroxylation of proline 70 on actin disrupts its interaction with SETD3, and suppresses actin methylation at histidine 73

Quantitative analysis of hydroxylated actin peptides revealed significant decrease of actin hydroxylation at Pro 70 (Fig. S2A). Interestingly, recent studies have demonstrated that SETD3 promotes actin polymerization by methylating a histidine residue at position 73^9^. Because of the spatial proximity of these two residues, we hypothesized that there could be a potential crosstalk between proline hydroxylation at position 70 and histidine methylation at position 73.

To test this hypothesis, we generated an actin mutant in which proline 70 was substituted with alanine (P70A). Immunoprecipitation assays showed that the interaction between the P70A mutant and SETD3 was significantly enhanced compared to wild-type actin (Fig. 3A), suggesting that hydroxylation of proline 70 may hinder SETD3 binding, thereby inhibiting histidine 73 methylation.

**Fig. 3.**
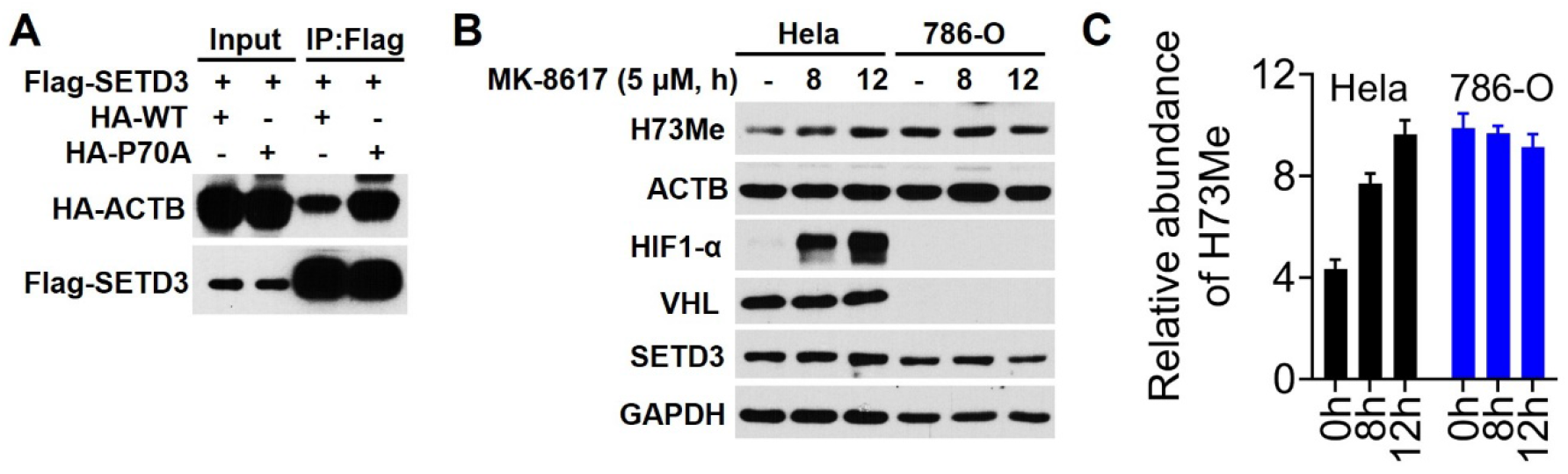
Hydroxylation of proline 70 on actin disrupts its interaction with SETD3 and inhibits its methylation at histidine 73. A. Pro70 mutation of actin blocks the interaction of SETD3 and actin. HA-tagged actin (wild type and P70A mutant) was co-transfected with Flag-tagged SETD3 into HEK293T cells. The cell lysates were subjected to immunoprecipitation using the anti-Flag beads. HA-actin (WT and P70A mutant) were probed using an antibody against the HA-tag. (B-C). MK-8617 increases H73Me-actin in Hela cells but not in 786-O cells. Hela and 786-O cells were treated with MK-8617 (5 μM) for indicated time points. Cell lysates were subjected to western blot (B) and H73Me level were quantified (C). GAPDH was used as the control.

Consistent with this hypothesis, western blot analysis revealed that MK-8617 treatment significantly increased histidine 73 methylation in VHL-expressing HeLa and 293T cells. Interestingly, in VHL-deficient 786-O and RCCJF cells, histidine 73 methylation on actin remained unchanged upon MK-8617 treatment (Fig. 3B, Fig. 3C, Fig. S2B, and Fig. S2C). Finally, we surveyed a panel of cell lines, and found that VHL-deficient cells are generally associated with higher actin histidine 73 methylation levels (Fig. S2D). These findings suggest that VHL may play a crucial role in modulating the crosstalk between proline hydroxylation at position 70 and histidine methylation at position 73 on actin.

### VHL inhibits SETD3-dependent methylation of histidine 73 on actin

To elucidate the mechanism by which VHL regulates the interplay between proline hydroxylation at position 70 and histidine methylation at position 73 on actin, we synthesized four biotinylated actin peptides. These peptides contained proline 70 in both hydroxylated and non-hydroxylated forms, with varying lengths (Fig. 4A). Biotin-streptavidin binding assays demonstrated that VHL selectively interacted with the hydroxylated, but not the non-hydroxylated peptides (Fig. 4B).

**Fig. 4.**
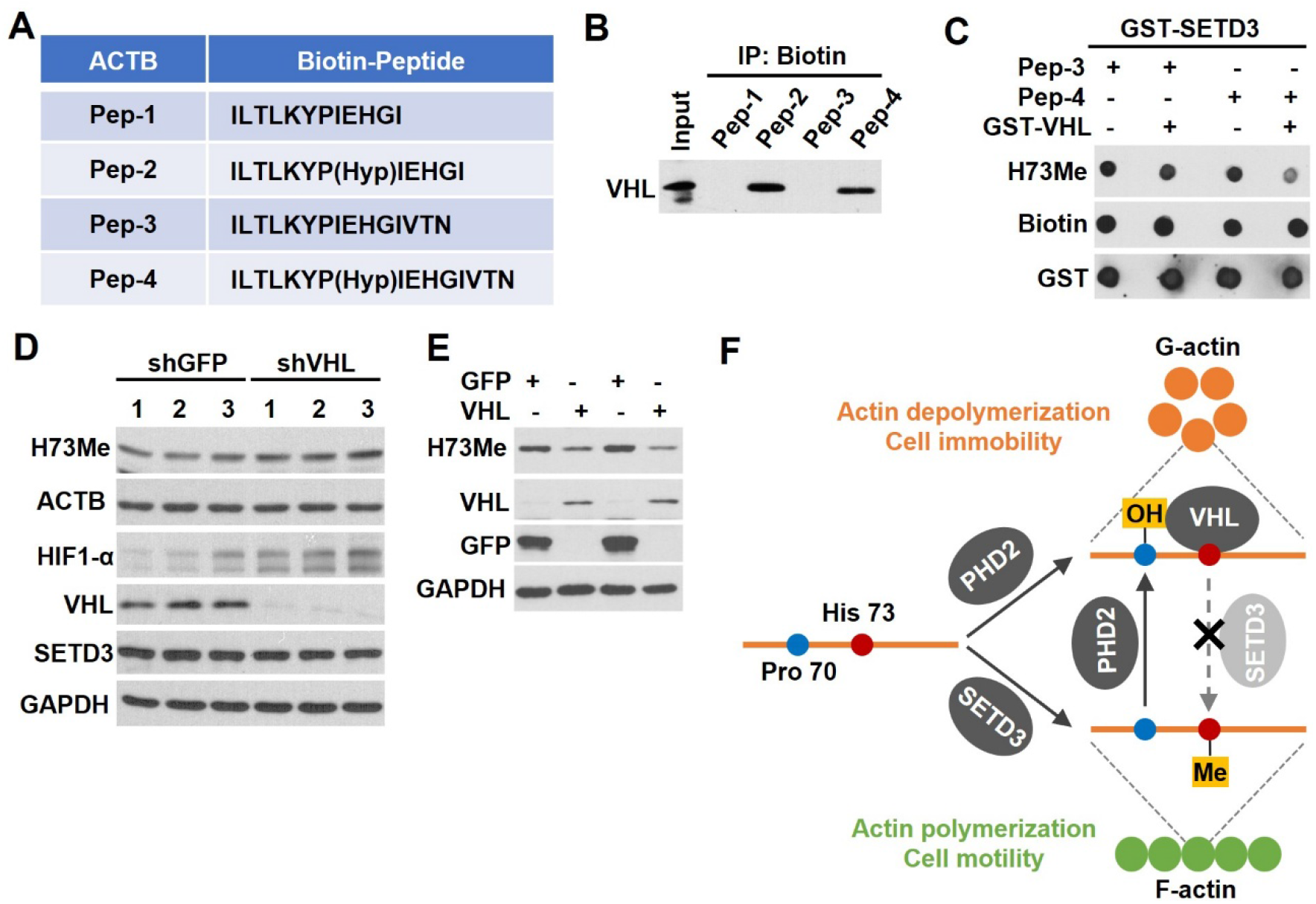
Hydroxylation of proline 70 on actin inhibits SETD3-mediated methylation of histidine 73 by recruiting VHL. A. The peptides sequence synthesized, including the non-hydroxylated and hydroxylated (Hyp70) version of the peptide. B. VHL interacts with the hydroxylated Pro70-actin peptide but not the non-hydroxylated version of the peptide. The peptides were incubated with the cell lysates from HEK293T cells, and were subjected to streptavidin pulldowns. C. VHL blocks H73 methylation in the hydroxylated (Pro70) peptide. In vitro methylation assay with synthesized WT and Hydroxylated (Pro70) peptides. D. VHL knockdown increases H73Me and HIF1a levels in Hela cells. Cell lysates were subjected to western blot. E. VHL overexpression decreases H73Me levels in 786-O cells. Cell lysates were subjected to western blot. F. A model of the mechanism by which Pro70 hydroxylation on actin regulates H73 methylation, actin polymerization, and cell motility.

Subsequently, in vitro methylation assays were performed using SETD3 and the hydroxylated peptides. The results showed that the presence of VHL significantly reduced SETD3-mediated methylation of histidine 73 on peptide 4 (containing hydroxylated Pro 70). On the contrary, the addition of the recombinant VHL protein did not affect SETD3-mediated methylation of histidine 73 on peptide 3 (non-hydroxylated) (Fig. 4C). Consistent with these findings, Western blot analysis showed a significant increase in histidine 73 methylation in HeLa cells upon VHL knockdown, whereas VHL overexpression markedly reduced histidine 73 methylation in 786-O cells (Fig. 4D, Fig. 4E, and Fig. S3A).

These results suggest that Pro 70 of actin is hydroxylated by the PHD enzymes, and this hydroxylation event is responsible for the recruitment of VHL. The binding of VHL to Pro-hydroxylated actin interferes with the binding and recruitment of SETD3, likely through steric hinderance. This interference reduces histidine 73 methylation, ultimately impairing actin polymerization and cell motility (Fig. 4F).

### Discussion PHDs function as a writer of proline hydroxylation on actin

Our hydroxylation proteomics analysis identified multiple proline hydroxylation sites on actin (Fig. 2H). Among these, Pro307 has been previously reported to be hydroxylated by PHD3, which inhibits actin filament formation and cell motility, but the precise molecular mechanism remains unclear^16^. Notably, quantitative proteomic analysis revealed that hydroxylation of Pro70 is suppressed by PHD inhibitor treatment (i.e., MK-8617) (Fig. 2C and Fig. 3A). Furthermore, we determined that PHD2 and PHD3, but not PHD1, interact with actin (Fig. 2A), suggesting that PHD2 or PHD3 mediate actin hydroxylation. These results provided evidence pointing to PHD2 and PHD3 as potential “writers” of actin hydroxylation. However, the distinct contributions of PHD2 and PHD3 to the hydroxylation of Pro70 on actin remain to be clarified. PHD2 is widely expressed across tissues, while PHD1 and PHD3 exhibit more restricted tissue expression patterns^17^.

Importantly, a missense mutation in ACTB (c.208C>G, p.Pro70Ala) has been identified in patients with Baraitser-Winter Syndrome (BWS)^18^, a genetic disorder that is associated with distinctive combinations of craniofacial, neurological, and developmental abnormalities. Additionally, amino acid sequence alignment reveals that Pro70 and His73 on actin are highly conserved across various actin isoforms and across multiple species (Fig.S3B and S3C). This data provide additional evidence pointing to the functional relevance of this PTM cross talk mechanism. At the same time, it is also important to recognize that these actin isoforms are not functionally redundant, with their specific roles determined by their unique expression patterns and interactions with various actin-binding proteins^19^. Consequently, the cross talk between Pro hydroxylation and His methylation on different actin isoforms is likely to result in diverse biological effects.

### VHL functions as a reader of proline hydroxylation on actin

We identified that hydroxylated actin is recognized and bound by VHL, suggesting that VHL is a “reader” of actin hydroxylation. This binding disrupts the interaction between actin and SETD3, thereby inhibiting the methylation of His73 on actin (Fig. 3B, Fig. 3C and Fig. 4C). The spatial proximity of Pro70 to His73 suggests that VHL could sterically block SETD3, preventing it from methylating His73. Interestingly, although VHL’s primary role as a component of the E3 ligase complex is to mediate the degradation of hydroxylated HIF-α under normoxic conditions^12^, hydroxylated actin recognized by VHL does not undergo this degradation pathway (Fig. 4B). Instead, VHL binds to hydroxylated actin, effectively competing with SETD3 and inhibiting its methylation activity. Consistently, ccRCC cell lines with inherited VHL loss-of-function did not respond to MK-8617 treatment, whereas reintroducing VHL significantly reduced histidine 73 methylation and cell mobility (Fig. 1A, Fig. 1C, Fig. 1D, and Fig. 4E). This suggests that actin hydroxylation by PHDs regulates actin function through a VHL-dependent mechanism, independent of the ubiquitin-proteasome pathway.

### Crosstalk between actin hydroxylation and methylation

Our data reveal that the mutually exclusive PHD-mediated hydroxylation and SETD3-mediated methylation constitute refined regulatory mechanisms for actin dynamics. This raises an important question: how do cells achieve a balance between actin hydroxylation and methylation? Notably, distinct from SETD3^9, 20^, PHD is an αKG-dependent enzyme whose activity is regulated by αKG, oxygen, and ferrous ions. This distinction implies that the availability of these cofactors may regulate the balance between actin hydroxylation and methylation. For instance, under normoxic conditions, active PHDs promote actin hydroxylation, suppressing actin methylation and cell motility. In contrast, hypoxic conditions inhibit PHD activity, which could then shift the balance toward actin methylation. This then could lead to enhanced cell motility, a mechanism likely contributing to cancer cell invasion and migration under hypoxic conditions. These insights highlight SETD3 as a potential target for therapeutically modulating the metastasis of VHL-proficient tumors (under hypoxic conditions), or VHL-deficient tumors (under either hypoxic or normoxic conditions).

The discovery of hypoxia-inducible factor (HIF)-prolyl hydroxylase inhibitors (PHIs) has revolutionized the therapeutic approach to anemia, particularly in patients with chronic kidney disease (CKD). However, studies suggest that PHD inhibition may drive tubulointerstitial fibrosis, which is a process that involves enhanced deposition of the extracellular matrix (ECM) and cell migration, with epithelial-mesenchymal transition (EMT) being pivotal^21^. During this process, tubular epithelial cells convert into myofibroblasts and move into the interstitium, promoting fibrosis. Our findings further indicate that PHD inhibition can enhance cell motility in a VHL-dependent manner. Therefore, careful consideration of potential adverse effects is essential when advancing PHD inhibitors into clinical applications.

In summary, we demonstrate that actin hydroxylated by PHD enzymes at Pro70. This modification events recruits the von Hippel-Lindau (VHL) tumor suppressor protein, which then disrupts the interaction between actin and SETD3, leading to reduced SETD3-mediated methylation of histidine 73 (His73) on actin is suppressed. This then results in impaired actin polymerization and compromised cell motility. Overall, this study not only expands the repertoire of actin PTMs, but also provides, for the first time, a mechanistic explanation of cytoskeletal dynamic regulation from the perspective of crosstalk between actin hydroxylation and methylation. Finally, SETD3-targeting inhibitors could have significant potential for modulating tumor invasion and metastasis, a critical aspect that warrants further research.

## Acknowledgments

This work was supported by grants from the NIH (R35GM134883)

---

**Fig. S1.**
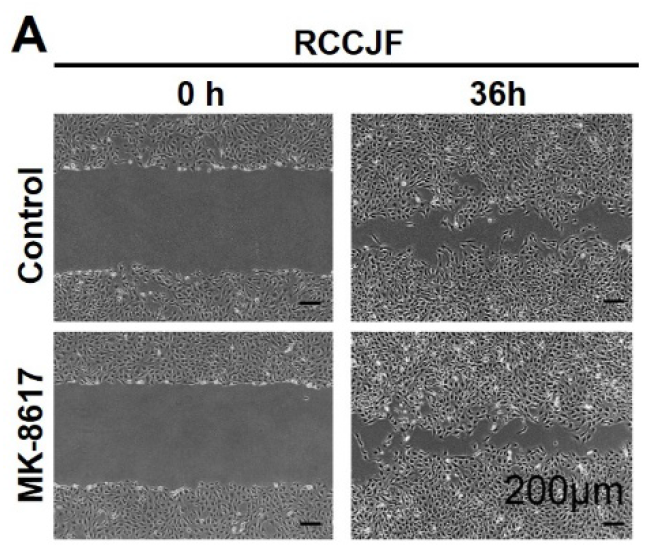
VHL deletion blocks the effect of MK-8617 in promoting cell motility. A. RCCJF cells were subjected to scratch assays with or without MK-8617 treatment (5 μM) for 36 hours. Representative images from two independent experiments were shown. Scale bar, 200 μm.

**Fig. S2.**
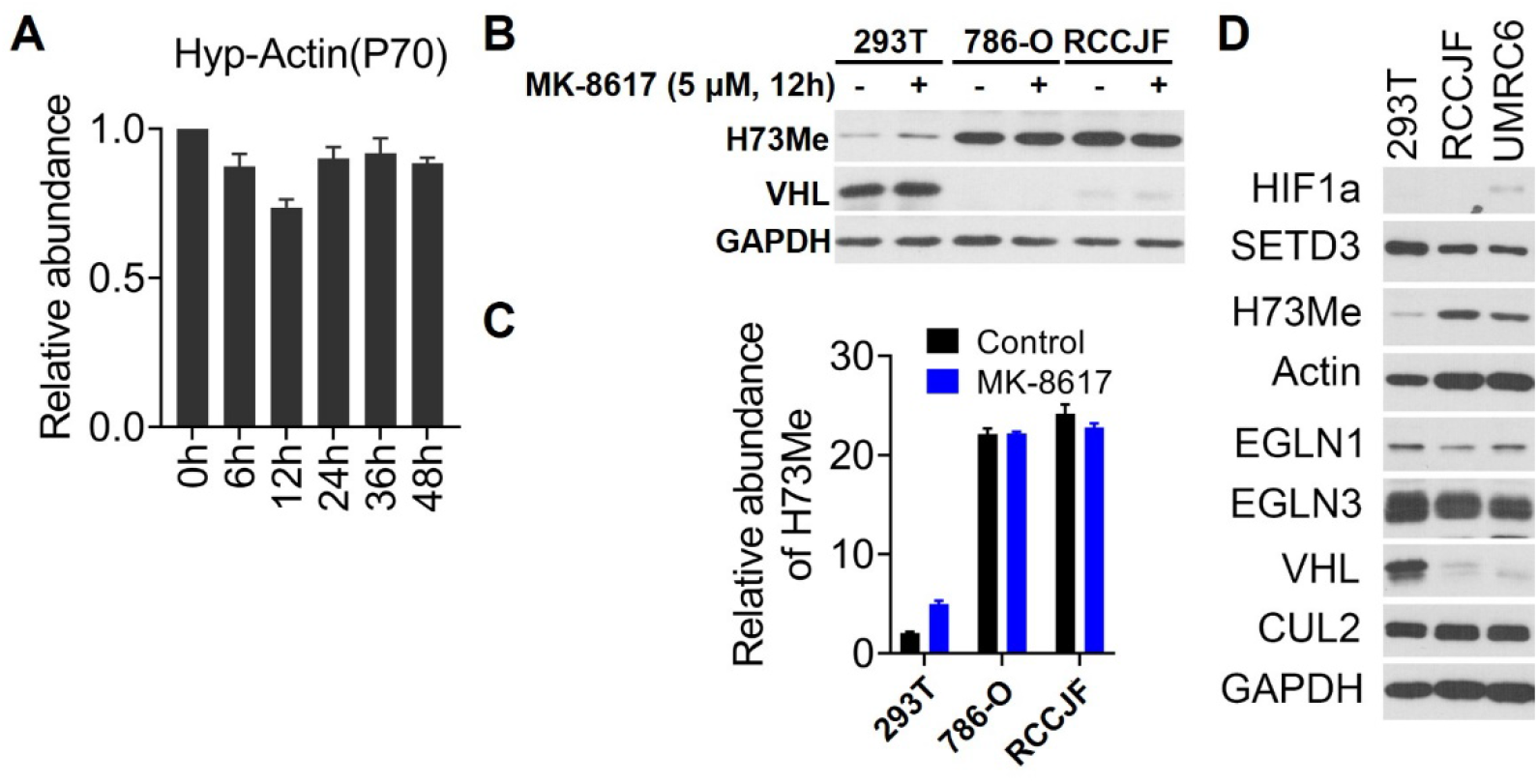
VHL-deficient cell lines exhibit significantly increased methylation of lysine 73 on actin. A. TMT-based quantitative proteomic analyses revealed the hydroxylation level of proline 70 on actin in MCF10A cells following MK-8617 treatment (5 μM) at the indicated time points. (B-C). MK-8617 increases H73Me-actin in 293T but not in 786-O and RCCJF cells. 293T, 786-O, and RCCJF cells were treated with MK-8617 (5 μM) for 12 hours. Cell lysates were subjected to western blot (B) and H73Me level were quantified (C). D. 293T, RCCJF, and UMRC6 cells were lysed and subjected to western blot analysis.

**Fig. S3.**
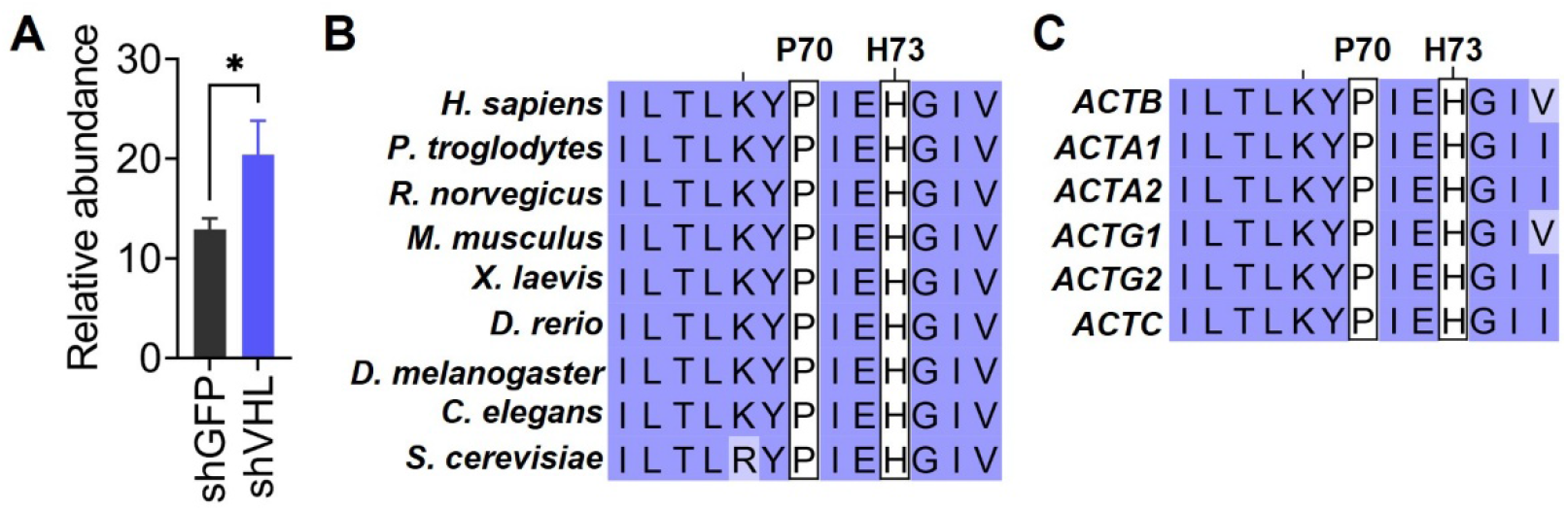
Proline 70 and Histidine 73 on actin are highly conserved across species and isoforms. A. VHL knockdown increases H73Me in Hela cells. Cell lysates were analyzed by western blot and H73Me levels were quantified. GAPDH was used as the control. Data are shown as mean ± SEM, n=3 each group. (B-C). Proline 70 and Histidine 73 on actin are highly conserved across different species and actin isoforms. Comparative analysis of sequences adjacent to amino acids 70 and 73 on ACTB Across nine species (B) and six actin isoforms (C).

